# Structural basis for no retinal binding in flotillin-associated rhodopsins

**DOI:** 10.1101/2025.04.29.651185

**Authors:** Kirill Kovalev, Artem Stetsenko, Florian Trunk, Egor Marin, Jose M Haro-Moreno, Gerrit H.U. Lamm, Alexey Alekseev, Francisco Rodriguez-Valera, Thomas R. Schneider, Josef Wachtveitl, Albert Guskov

## Abstract

Rhodopsins are light-sensitive membrane proteins capturing solar energy via a retinal cofactor covalently attached to a lysine residue. Several groups of rhodopsins were reported to lack the conserved lysine and showed no retinal binding. Recently, flotillin-associated rhodopsins (FArhodopsins) were identified and suggested to lack the typical retinal binding pocket despite preserving the lysine residue in many members of the group. Here we present cryo-EM structures of paralog FArhodopsin and proteorhodopsin from marine bacteria *Pseudothioglobus*. The structures revealed pentameric assemblies of both proteins similar to those of other microbial rhodopsins. We demonstrate no binding of retinal to the FArhodopsin despite preservation of the lysine residue and overall similarity of the protein fold and internal organization to those of the retinal-binding paralog. Mutational analysis confirmed that two amino acids, H84 and E120, prevent retinal binding within the FArhodopsin. Thus, our work provides insights into the natural retinal loss in microbial rhodopsins and might contribute to the further understanding of the FArhodopsin clade.

## INTRODUCTION

Microbial rhodopsins (MRs) are light-sensitive membrane proteins capable of performing various functions, including ion transport, signal transduction, or enzymatic activity, and play vital roles in bioenergetics and phototaxis of numerous microorganisms^1^. Despite such a great diversity, most MRs have a core of 7 transmembrane (TM) helices, typically named A to G, and sense visual light via the retinal cofactor, covalently attached to the highly-conserved lysine residue in the 7th TM helix (G).

However, there are many natural exceptions lacking this lysine residue (Rh-noK)^2^. For instance, Rh-noK have been described for a group of xanthorhodopsin-related sequences in marine betaproteobacteria and were shown to be retinal binding and functional proton pumps when the lysine residue was reconstituted by genetic manipulation^3^. Interestingly, Rh-noK was found located in tandem with a functional proteorhodopsin (PR) gene, and the authors speculated that both gene products could form a multimer in which Rh-noK could regulate the proton pumping activity of the neighbor^3^.

Another example is a clade of recently found flotillin-associated rhodopsins (FArhodopsins or FARs), which were suggested to lack retinal cofactor not due to absent lysine, but because of a reduced retinal-binding pocket (RBP)^4^. Indeed, marine FARs maintain the retinal-binding lysine residue in the 7th TM helix, which is in contrast to Rh-noK. Both FARs and Rh-noK were found in genomes containing also a paralog rhodopsin gene that were predicted to encode for standard proton-pumping PRs. Rhodopsin paralogs (homologous genes found within the same genome) are widespread in haloarchaea where they have been extensively studied and revealed to have often different functions^5,6^. On the other hand, marine bacteria with streamlined genomes rarely have paralogs^7,8^. Thus, the finding of proton-pumping paralogs in the two known cases of retinal-less rhodopsins points to radically novel functions for them.

Given that all microbes producing FARs belong to uncultivated aquatic microbes, we have resorted to expression in a surrogate host. We cloned and expressed in *E*.*coli* two paralog rhodopsins found in a single amplified genome (SAG) classified as gammaproteobacterium *Pseudothioglobus sp*. To reveal whether marine FARs are capable of binding retinal and to gain insights into the structural organization of FArhodopsins, we used single-particle cryo-electron microscopy (cryo-EM) and obtained high-resolution structures of paralog PR and FAR from the marine SAG AG-319-B05 *Pseudothioglobus* (*Ps*PR and *Ps*FAR, respectively) allowing for direct comparison of the two types of rhodopsins found within a single genome. Our data demonstrates that, as expected, the architecture of *Ps*FAR is similar to those of the *Ps*PR and other MRs. However, we observed that, and as predicted in Haro-Moreno et al^4^, there was no retinal binding in the *Ps*FAR due to a dramatic decrease of its RBP, which is occupied by the side chains of the H84 and E120 residues. We demonstrate that replacement of both of these residues with more compact ones allows recovery of an effective retinal binding to *Ps*FAR containing lysine at residue K213 with only minor changes in the structure of the protein. At the same time, single mutations were not sufficient for that purpose.

## RESULTS

### Functional and structural characterization of *Ps*PR and *Ps*FAR

We produced *Ps*PR and *Ps*FAR as described elsewhere^9^. In line with predictions made in Haro-Moreno et al^4^, we found no signs of retinal binding to *Ps*FAR, which appeared colorless despite being supplemented with sufficient amounts of all-*trans* retinal during all stages of expression and purification (Fig. 1A). On the contrary, *Ps*PR demonstrated UV/Vis absorption spectra typical to that of normal rhodopsins^10^ with a maximum absorption wavelength of 512 nm (Fig. 1A). Spectral properties of *Ps*PR are similar to those of PRs with the pK_a_ of the retinal Schiff base (RSB) counterion of ∼5.1^11^ (Fig. 1B,C). The photocycle kinetics of *Ps*PR also resembles those of PRs^12^ (Fig. 1D,E; Fig. S1). *Ps*PR pumps protons as demonstrated by the pH changes in *E*.*coli* cell suspension (Fig. 1F). In summary, *Ps*PR represents a typical green-light-absorbing proteorhodopsin (GPR).

**Fig. 1.**
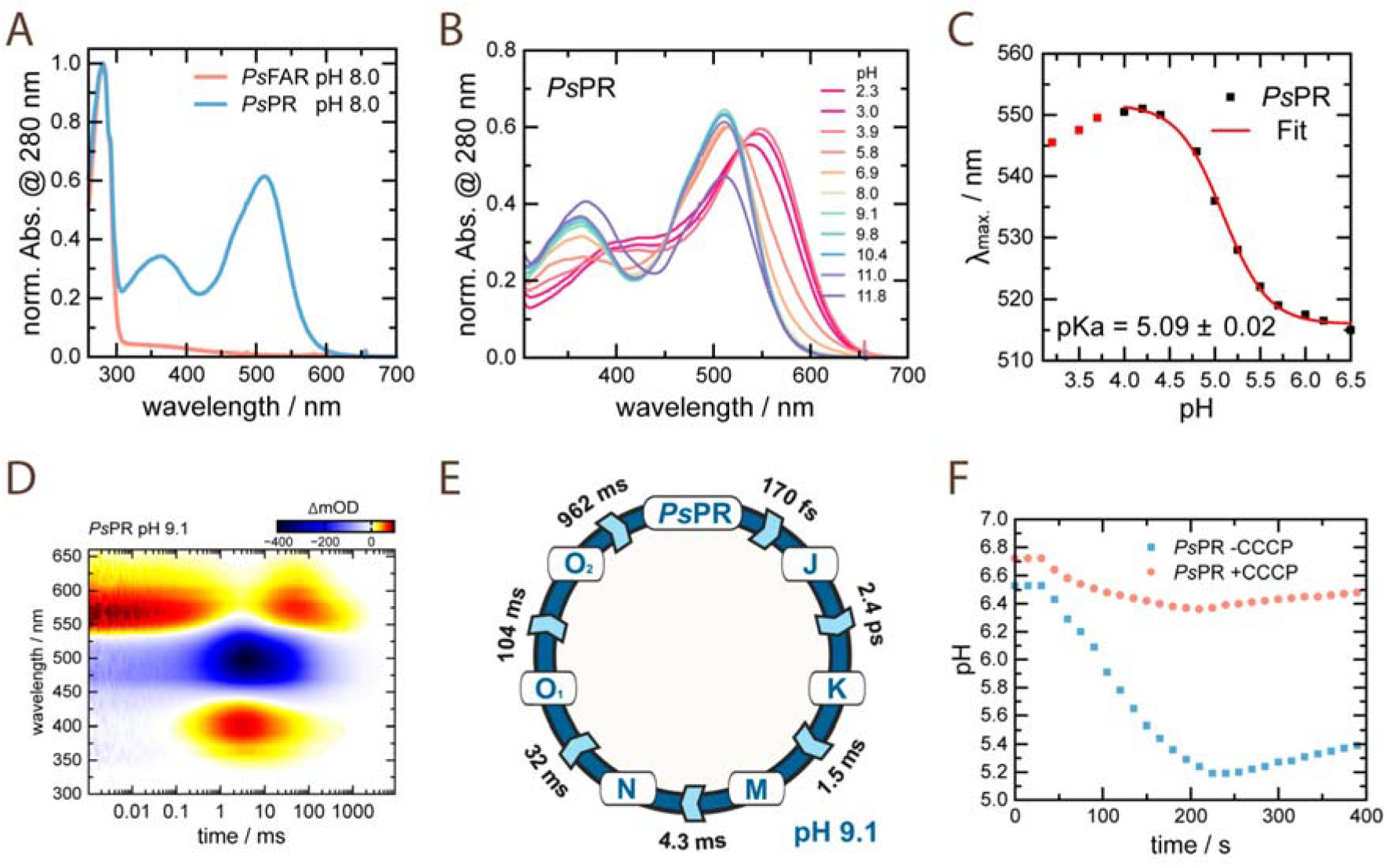
Spectroscopic characterization of *Ps*PR and *Ps*FAR. **A**. Absorption spectra of purified *Ps*PR and *Ps*FAR at pH 8.0. **B**. pH titration of *Ps*PR. The spectra are normalized to the absorption at 280 nm. **C**. Determination of pKa value of *Ps*PR. **D**. Photocycle kinetics depicted as 2D-contour plot. The signal amplitude is color coded as follows: positive (red), no (white) and negative (blue) absorbance changes **E**. Scheme of the proposed *Ps*PR photocycle. **F**. Proton pumping activity tests of *Ps*PR in *E*.*coli* cell suspension.

Next, we used single-particle cryo-EM and obtained 2.5Å-resolution structures of the *Ps*PR and *Ps*FAR proteins (Fig. S2; Table S1). Direct comparison of the two rhodopsins from the same genome (SAG) sharing >62% sequence similarly (>44% sequence identity) (Fig. S3) helped us to establish functional differences carried out in the same cell. Furthermore, we also compared the structure of *Ps*FAR to those of other known MRs to better understand its peculiarities.

### Absence of retinal binding in *Ps*FAR

As evidenced by the absence of typical retinal-associated absorption maxima in the *Ps*PAR spectrum, we found no retinal cofactor bound to the FAR in the structure (Fig. 2A). While the lysine residue in helix G is preserved (K213) and is oriented similarly to the respective residue of *Ps*PR (K214), there is no space to allow retinal binding in the central region of the rhodopsin (Fig. 2A,B).

**Fig. 2.**
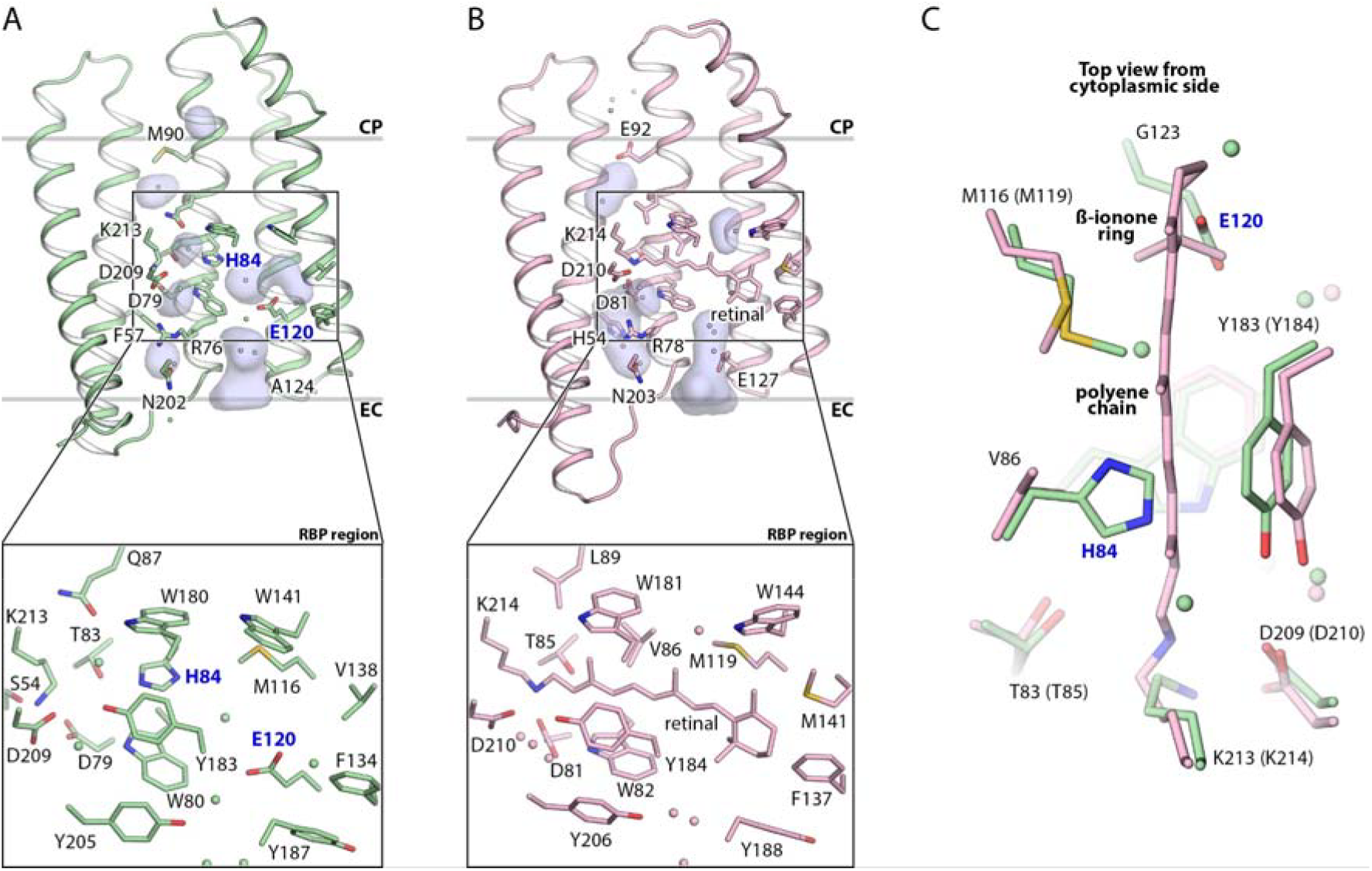
Region of retinal binding pocket in *Ps*FAR and *Ps*PR. **A**. Side view of the *Ps*FAR protomer (top panel) and detailed view of the RBP region (bottom panel). H84 and E120 substituting for retinal cofactor are indicated in blue. **B**. Side view of the *Ps*PR protomer (top panel) and detailed view of the RBP region (bottom panel). Internal cavities were calculated using HOLLOW^14^ and are shown with light blue surfaces. **C**. Top view from the cytoplasmic side onto the alignment of the RBP regions of *Ps*FAR (green) and *Ps*PR (pink).

Firstly, the central part where the polyene chain of retinal is expected is occupied by the side chain of H84 (Fig. 2C). Secondly, the region of the ß-ionone ring is largely occupied by the side chain of E120 (Fig. 2C). This observation is in line with the prediction made in Haro-Moreno et al^4^. The presence of H84 is likely the major reason for no retinal binding in *Ps*FAR as it results in only fragmented cavities of small size instead of continuous retinal binding pocket found in *Ps*PR and other MRs (Fig. 2A; Fig. S4). Moreover, aromatic amino acid residues, such as histidine or phenylalanine, are found in all FARs, while E120 is less conserved within the clade (Fig. S5). For instance, it is replaced with small glycine in the subgroup HIMB59; however, in this case, tryptophan is found at the position of G137 of *Ps*FAR, likely occupying the space of the ß-ionone ring of the retinal (Fig. S5). As described further in the manuscript, both H84 and E120 play key roles in blocking the RBP of *Ps*FAR.

The internal cavities in the central region of the *Ps*FAR protomer are occupied with water molecules, mediating a hydrogen bond network likely stabilizing the protein (Fig. S4). The resolution of the *Ps*FAR structure does not allow to univocally identify the interaction partner of K213; nevertheless, the maps suggest that it is stabilized by H-bonds to the S54, D79, and D209 residues. Analysis of the sequences of paralog FARs and PRs from different organisms showed that S54 is conserved in FArhodopsins but is replaced with alanine in PRs (Fig. S5). This additionally highlights the importance of S54 for stabilization of lysine residue not connected to a cofactor in FARs.

Noteworthy, although the arrangement of TM helices of *Ps*PAR in the central region is similar to that of *Ps*PR, there is a 2Å-shift of helix E of *Ps*FAR towards the center of the protomer further reducing the space for the ß-ionone ring of the retinal cofactor (Fig. S6).

### Oligomerization of FArhodopsin

Both *Ps*FAR and *Ps*PR form pentamers of similar type, resembling those of other pentameric MRs, such as PRs^15,16^, light-driven sodium pumps (NaRs)^9,17,18^, or viral rhodopsins of group 2^19^ (Fig. 3). The architecture of the protomers is also very similar in *Ps*PR and *Ps*FAR. In the FAR, helix G is 3 turns longer than that of *Ps*PR and sticks out towards the cytoplasmic side by an additional 12Å, similar to what was shown for BPR. This results in the more prominent cytoplasmic part of the *Ps*FAR pentamer (Fig. 3). On the contrary, the extracellular side is larger in *Ps*PR due to the 1 turn longer helix B and elongated BC-loop. In general, *Ps*PR has a similar pentamer profile as that of GPR while *Ps*FAR is closer to BPR (Fig. 3).

**Fig. 3.**
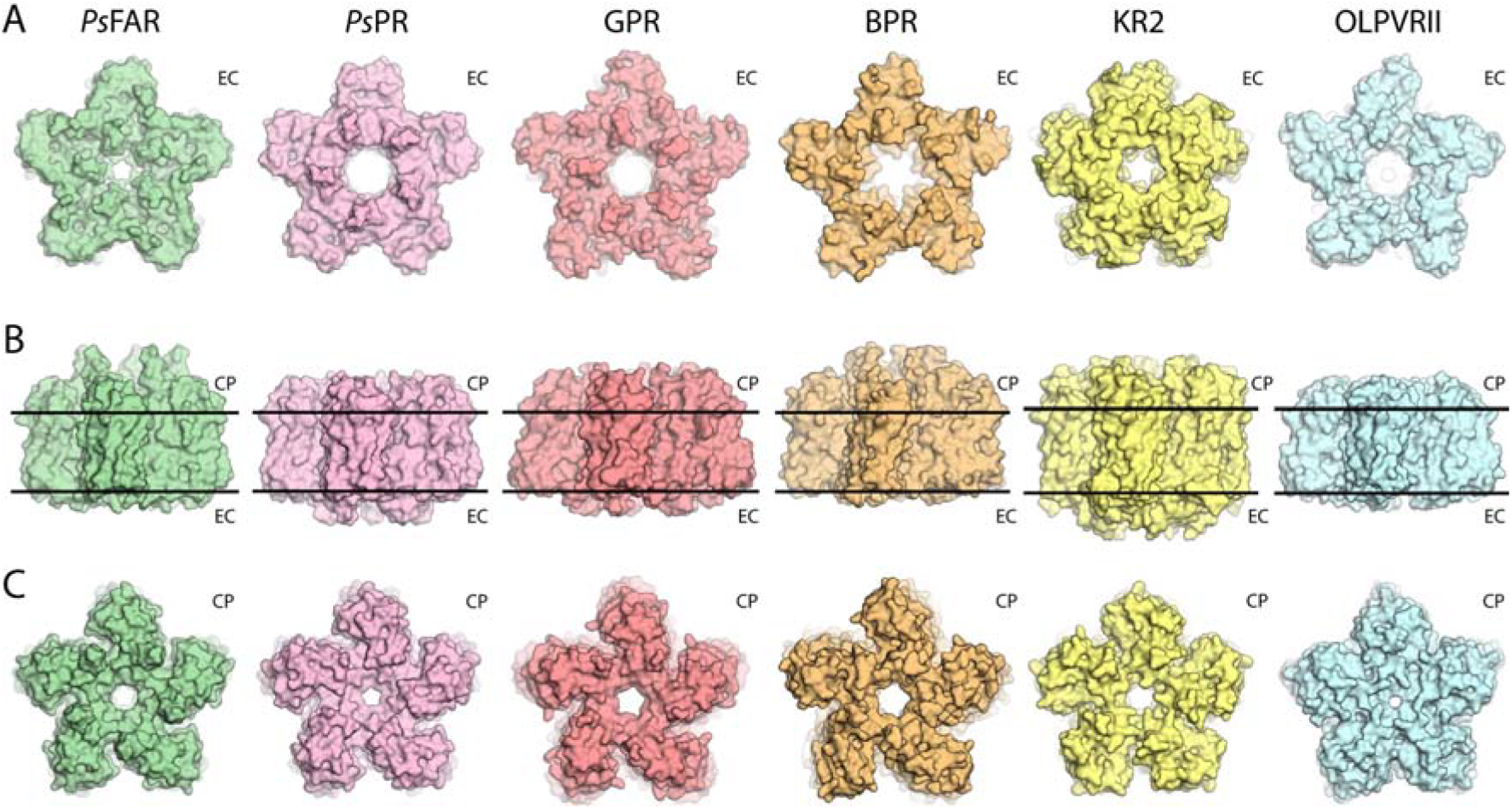
Oligomers of *Ps*FAR and *Ps*PR and comparison to other MRs. **A**. View from the extracellular side. **B**. Side view of the oligomers in the membrane. The hydrophobic/hydrophilic membrane core boundaries have been calculated using PPM server^20^ and are shown with black lines. **C**. View from the cytoplasmic side. *Ps*FAR (present work) is colored green. *Ps*PR (present work) is colored pink. Green-light-absorbing PR (GPR, PDB ID: 7B03^16^) is colored red. Blue-light-absorbing PR (BPR, PDB ID: 4KLY^15^) is colored orange. Light-driven sodium pump KR2 (PDB ID: 6YC3^21^) is colored yellow. Viral rhodopsin of group 2 (OLPVRII, PDB ID: 6SQG^19^) is colored cyan.

The interprotomeric contacts differ in *Ps*FAR and *Ps*PR (Fig. S7). First, helix B of *Ps*FAR is shifted towards the center of the pentamer compared to that in *Ps*PR, with the maximal amplitude of nearly 5Å at its extracellular tip (Fig. S7A). Second, the side chain of Y59 in *Ps*FAR is oriented towards the center of the pentamer compared to the side chain of corresponding residue Y56 in *Ps*PR (Fig. S7B,C). This reorientation results in a much smaller diameter of the central region of the *Ps*FAR oligomer (10 Å in *Ps*FAR vs. 20 Å in *Ps*PR) (Fig. S7B,C).

The difference in the Y59 and Y56 positions in *Ps*FAR and *Ps*PR, respectively, likely originates from the absence in *Ps*FAR of the histidine residue in helix B (H54 in *Ps*PR), which is conserved within PRs and plays an important role interacting with primary proton acceptor while being also involved in the interprotomeric contacts^22^ (Fig. S7D,E). For example, in *Ps*PR, H54 interacts with W13 and R59 of the nearby protomer via a water molecule (Fig. S7E). In *Ps*FAR, the histidine is replaced with phenylalanine (F57), and the interprotomeric contacts are organized mostly via the Y60’-S19 hydrogen bond (Fig. S7D). As a result, the side chain of Y60 leaves no space for Y59, which is consequently oriented towards the center of the pentamer.

### Conservation of the typical rhodopsin architecture in *Ps*FAR

Not only the positions of side chains of K213 and K214 are similar in *Ps*FAR and *Ps*PR, respectively, but the overall internal organization is well-preserved in the FAR despite the absence of the retinal cofactor. The central regions with the lysine residue and carboxylic counterions of the RSB of typical rhodopsin appear similar in *Ps*FAR and *Ps*PR. However, the side chains of D79 and D81 are oriented slightly differently. Also, as mentioned above, the conserved histidine residue in PRs, H54 in *Ps*PR, is absent in *Ps*FAR and is substituted with phenylalanine F57.

The extracellular (EC) internal parts of both rhodopsins between the arginine (R76 in *Ps*FAR and R78 in *Ps*PR) and the bulk are very similar. However, while there are rechargeable residues D72 and E127 at the EC part of *Ps*PR, which might be involved in proton transport, *Ps*FAR lacks carboxylic residues in the corresponding region.

Finally, the cytoplasmic (CP) parts of *Ps*PR and *Ps*FAR are also similar and mostly hydrophobic. *Ps*PR has a proton donor residue E92, which is absent in *Ps*FAR and replaced by methionine (M90).

In summary, the architecture of the *Ps*FAR protomer is very similar to that of *Ps*PR and many other MRs. This observation is in line with predictions made in Haro-Moreno et al^4^, and also is similar to what was suggested for Rh-noK^3^.

### Recovery of *Ps*FAR retinal binding

To probe whether H84 and E120 are the only determinants of no retinal binding to *Ps*FAR we mutated these residues to those found at the corresponding positions of *Ps*PR. Namely, we tested retinal binding to the *Ps*FAR-H84V, *Ps*FAR-E120G, and *Ps*FAR-H84V/E120G (hereafter *Ps*FAR-DM) variants. Expression and purification of the mutants supplemented with excess amounts of all-*trans* retinal demonstrated that only the replacement of both amino acid side chains results in effective retinal binding, while single mutations were not sufficient for recovery of the cofactor binding (Fig. 4A).

**Fig. 4.**
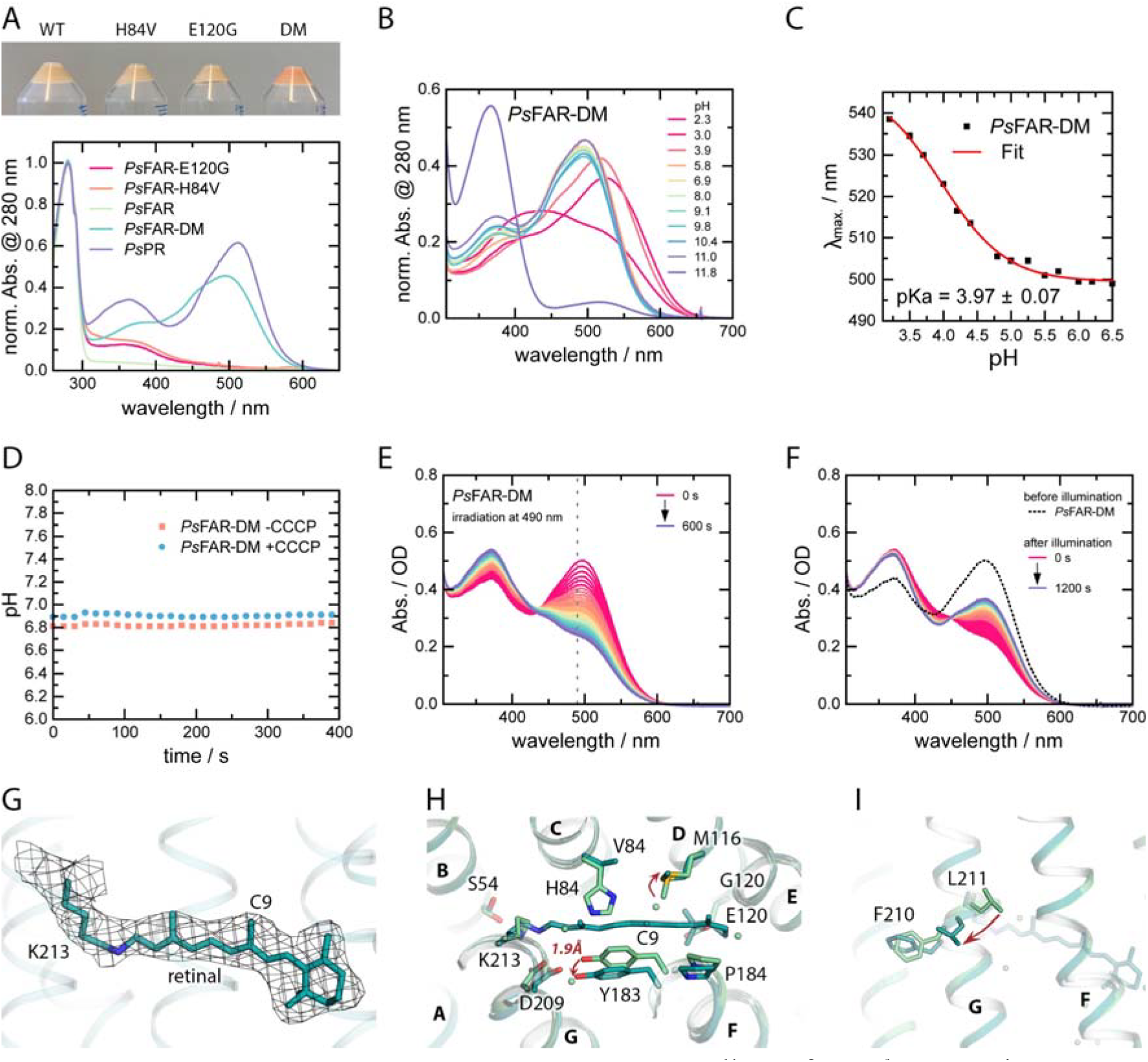
Recovery of retinal binding to *Ps*FAR. **A**. Pellets of *E*.*coli* expressing *Ps*FAR variants (top) and corresponding absorption spectra of the purified rhodopsins (bottom). **B**. pH titration of the absorption spectra of *Ps*FAR-DM. **C**. pK_a_ value determination of the RSB counterion at acidic pH. **D**. Ion pumping activity tests of *Ps*FAR-DM in *E*.*coli* cell suspension. **E**. Evolution of the absorption spectra of purified *Ps*FAR-DM upon continuous illumination with 490 nm LED. **F**. Recovery of the absorption spectra of purified *Ps*FAR-DM after continuous illumination with 490 nm LED. **G**. All-*trans* retinal in the cryo-EM structure of *Ps*FAR-DM covalently attached to K213. **H**. Alignment of *Ps*FAR-WT (green) and *Ps*FAR-DM (cyan). Red arrows indicate the main changes in the structure. Helices are indicated with bold capital letters (A-G). **I**. Rearrangements of helix G upon retinal binding to *Ps*FAR-DM. Red arrow indicates the flip of L211 towards the adjacent protomer in *Ps*FAR-DM compared to *Ps*FAR-WT.

At neutral pH, *Ps*FAR-DM demonstrated an absorption maximum at 498 nm, closer to that of BPR^23^, in line with overall structure similarities of BPR and *Ps*FAR (Fig. 1A, 4A). Also, the presence of glutamine in helix C (Q87) of *Ps*FAR is known as a signature of PRs with a blue-shifted spectral peak^24^. The pKa of the RSB counterion is lower than that of *Ps*PR and is ∼4 (Fig. 4B,C). *Ps*FAR-DM demonstrated no proton transport activity, which is in line at least with the absence of the proton donor residue (M90) in the FAR (Fig. 4D).

To probe whether *Ps*FAR-DM has a typical rhodopsin photodynamics upon light activation, we used femtosecond transient absorption spectroscopy (Fig. S8). Unlike *Ps*PR, which demonstrated similar behavior to that of the H75N mutant of GPR with deprotonated RSB counterion D97^12,25^ (Fig. S8A), *Ps*FAR-DM showed unusual ultrafast kinetics. Specifically, in *Ps*FAR-DM, the J state is missing, while the signatures of the excited state absorption (ESA) and stimulated emission (SE) centered around 540 nm and >640 nm, respectively, are present up to 10 ps. This is in contrast to *Ps*PR, where both ESA and SE disappear after only 200 fs. Overall, the ultrafast kinetics of *Ps*FAR-DM is reminiscent of that of some MRs with protonated RSB counterion^26,27^. Nevertheless, in both *Ps*PR and *Ps*FAR-DM, the non-decaying K or K-like intermediate state is formed within 2.4 ps and >10 ps, respectively (Fig. S8). These intermediates are spectrally similar, and are assumed to contain the isomerized chromophore. It is apparent that *Ps*FAR-DM is less efficient than *Ps*PR, while still showing the expected rhodopsin photodynamics on this timescale^28^.

Illumination experiments of *Ps*FAR-DM revealed a slow photocycle kinetics, resulting in almost bi-stable behavior of the protein. Specifically, upon illumination with 490 nm LED light, the photostationary state (PSS) at 375 nm is accumulated (Fig. 4E), which only partially and slowly relaxes back to the 490 nm-absorbing-state over time after switching off the illumination (Fig. 4F). The PSS at 375 nm likely reflects the photocycle intermediate with deprotonated RSB, known as M-state. This is again in line with the absence of the proton donor at the cytoplasmic side, hampering the reprotonation of the RSB in *Ps*FAR-DM.

To further understand whether the recovered binding of the retinal in *Ps*FAR-DM resembles that in *Ps*PR and other MRs, we obtained cryo-EM structure of the mutant at 2.8Å resolution. The structure of *Ps*FAR-DM is highly similar to that of the WT protein (root mean square deviation of 0.2 Å); however, it demonstrated an all-*trans* retinal covalently attached to K213 (Fig. 4G; Fig. S4). Upon cofactor binding, the residues in the retinal binding pocket are slightly rearranged. Most significantly, Y183 and P184 of helix F are shifted by ∼1.9Å to allow enough space for the polyene chain and ß-ionone ring, respectively (Fig. 4H). M116 is reoriented to avoid steric conflict with the C9 methyl group of the cofactor (Fig. 4H). Interestingly, upon retinal binding to K213, some other residues in helix G are rearranged. For instance, L211 is flipped by ∼5Å towards the adjacent protomer (Fig. 4I). There are no other notable changes either on the protein surface or inside the protomers, further demonstrating that the classical rhodopsin fold is preserved regardless of the retinal cofactor binding in the FARhodopsin.

## DISCUSSION

Here we presented the high-resolution structure of a flotillin-associated rhodopsin lacking the retinal cofactor. Direct comparison with the structure of a paralog proteorhodopsin located in the same genome (a marine SAG) allowed us to conclude that two amino acid residues, H84 and E120, play a key role resulting in no retinal binding in the marine FAR. Spectroscopic studies of the mutants with H84 and/or E120 substituted by more compact residues showed that not single, but only double mutation results in recovery of the retinal binding. The side chain of H84 is well-stabilized by H-bonds and replaces the central part of the polyene chain of retinal, leaving no space for the cofactor; additionally, aromatic residues at the position of H84 are conserved within FARs. Thus, the critical role of H84 can be expected. The absence of space for the retinal polyene chain in *Ps*FAR strongly suggests that the cofactor cannot bind to the rhodopsin not only in the tested all-*trans* configuration, but also in other conformations. On the contrary, E120 is flexible as evidenced by fragmented density maps around its side chain, and has several surrounding cavities; therefore, it might be in principle reoriented and allow enough space for the ß-ionone ring. Also, E120 is less conserved within the clade. Therefore, the necessity of the E120 replacement for the retinal binding is less expected and intriguing.

To the best of our knowledge the structure of *Ps*FAR is the first one of a microbial rhodopsin without retinal. It demonstrates that folding, internal organisation, and oligomeric assembly of FAR are preserved despite the absence of the cofactor and provides insights into the principles of this preservation.

Lastly, we found no obvious evidence of the FAR modification towards the interaction with flotillin compared to the PR. The elongation of the 7th TM helix towards the cytoplasm can be considered as one of the sites for protein-protein interactions (PPIs). Conformational changes in the protein surface found exclusively in the central part of helix G found upon the retinal binding recovery demonstrate flexibility of this region and might also suggest that helix G is involved in PPIs with flotillin. Importantly, helix G in the pentameric assembly is accessible for the PPIs within the lipid membrane.

Another region that is interesting in this aspect is the central part of helix E, located on the outside of the protein pentamer and being displaced in the FAR compared to the PR. Thus, one can speculate that helix E might be involved in the PPIs with flotillin; however, both hypotheses should be tested in future. In summary, the structure of *Ps*FAR shows conservation of the classical rhodopsin fold in the absence of retinal cofactor and highlights the importance of further studies of the intact rhodopsin-flotillin complexes to understand the biological role and function of FARs.

## RESOURCE AVAILABILITY

### Lead contact

Further information and requests for resources and reagents should be directed to and will be fulfilled by the lead contacts, Kirill Kovalev and Albert Guskov.

### Data and code availability

All atomic models and density maps presented in this study are publicly available as of the date of publication at the Protein DataBank (PDB IDs: 9R21, 9R22, and 9R23) and Electron Microscopy DataBank (EMDB IDs: EMD-53519, EMD-53520, and EMD-53521) for the structures of *Ps*FAR-WT, *Ps*PR, and *Ps*FAR-DM, respectively.

This paper does not report original code.

Any additional information required to reanalyze the data reported in this paper is available from the lead contacts upon request.

## ACKNOWLEDGEMENTS

E.M., A.S. and A.G. thank NeCEN personnel for the support during data collection. The access to NeCEN facilities was funded by the Netherlands Electron Microscopy Infrastructure (NEMI), project number 184.034.014 of the National Roadmap for Large-Scale Research Infrastructure of the Dutch Research Council (NWO). A.G. was supported by NWO grant OCENW.KLEIN.141. K.K. has been supported by EMBL Interdisciplinary Postdoctoral Fellowship (EIPOD4) under Marie Sklodowska-Curie Actions Cofund (grant agreement number 847543). The work of K.K. and T.R.S. was supported by the European Molecular Biology Laboratory. This research was supported by the German Research Foundation (CRC 1507 – Membrane-associated Protein Assemblies, Machineries and Supercomplexes; Project 05 to J.W.).

## AUTHOR CONTRIBUTIONS

K.K. did cloning, protein production, and functional tests in *E*.*coli* cell suspension; A.S. prepared grids for cryo-EM; E.M. and A.S. processed cryo-EM data; K.K. refined the cryo-EM structures; A.A. helped with the analysis of the structures; A.G. supervised all steps of cryo-EM pipeline and structure refinement and validation; F.T. performed spectroscopy studies of the purified proteins; G.H.U.L. helped with spectroscopy data analysis; J.W. supervised spectroscopy studies; J.M.H.-M. and F.R.-V. provided nucleotide and amino acid sequences of *Ps*FAR and *Ps*PR; J.M.H.-M. performed sequence alignments and prepared respective figures; K.K. prepared initial draft of the manuscript; all authors contributed to the preparation of the final manuscript.

## METHODS

### Cloning

The DNA coding *Ps*PR and *Ps*FAR were optimized for *E*.*coli* using GeneArt (Thermo Fisher Scientific). Genes were synthesized commercially (IDTDNA). For protein expression and purification, the pET15b plasmid with 6xHis-tag at the C-terminal was used. Point mutations were introduced using whole-plasmid PCR followed by the blunt ends ligation.

### Protein expression, solubilization, and purification

*E*.*coli* cells were transformed with pET15b plasmid containing the gene of interest. Transformed cells were grown at 37°C in shaking baffled flasks in an autoinducing medium ZYP-5052^29^, containing 10 mg/L ampicillin. They were induced at an OD_600_ of 0.8-0.9 with 1 mM isopropyl-β-D-thiogalactopyranoside (IPTG). Subsequently, 10 μM all-*trans*-retinal was added. Incubation continued for 3 hours. The cells were collected by centrifugation at 4000*g* for 25 min. Collected cells were disrupted in an M-110P Lab Homogenizer (Microfluidics) at 25,000 p.s.i. in a buffer containing 20 mM Tris-HCl, pH 8.0, 5% glycerol, 0.5% Triton X-100 (Sigma-Aldrich) and 50 mg/L DNase I (Sigma-Aldrich). The membrane fraction of the cell lysate was isolated by ultracentrifugation at 125,000*g* for 1 h at 4°C. The pellet was resuspended in a buffer containing 20 mM Tris-HCl, pH 8.0, 0.2 M NaCl and 1% DDM (Anatrace, Affymetrix) and stirred overnight for solubilization. The insoluble fraction was removed by ultracentrifugation at 125,000*g* for 1h at 4 °C. The supernatant was loaded on a Ni-NTA column (Qiagen), and the protein was eluted in a buffer containing 20 mM Tris-HCl, pH 8.0, 0.2 M NaCl, 0.4 M imidazole, and 0.1% DDM. The eluate was subjected to size-exclusion chromatography on a Superdex 200i 300/10 (GE Healthcare Life Sciences) in a buffer containing 20 mM Tris-HCl, pH 8.0, 100 mM NaCl and 0.04% DDM. In the end, protein was concentrated to 60 mg/ml and kept in –80°C for storage.

### Measurements of pump activity in the *E. coli* cells suspension

*Ps*PR, *Ps*FAR, and *Ps*FAR-DM were expressed as described above. The cells were collected by centrifugation at 4,000*g* for 15□min and were washed three times with an unbuffered 100 mM NaCl solution, with 30-min intervals between the washes to allow for exchange of the ions inside the cells with the bulk. After that, the cells were resuspended in an unbuffered 100 mM NaCl solution and adjusted to an OD_600_ of 8.5. The measurements were performed in 3-ml aliquots of stirred cell suspension kept at ice-cold temperature (0.3–1.0□°C). For measurements at higher temperature, the cell suspension were kept at room temperature (20– 22□°C). The cells were illuminated using a halogen lamp, and the light-induced pH changes were monitored with a pH meter (Mettler Toledo). The measurements were repeated under the same conditions after the addition of 30□μM CCCP protonophore. For the measurements of the pH changes at low pH values, the pH of the final cell suspension was adjusted using 10% HCl solution.

### Steady-state absorption spectroscopy, pH titration and illumination experiments

Stationary absorption spectra of investigated samples were obtained by a Specord S600 absorption spectrometer (Analytic Jena) using an optical path length of 10 mm. For all measurements, small amounts of protein stock were diluted in a buffer containing 10 mM sodium citrate, 10 mM MES, 10 mM HEPES, 10 mM Tris, 10 mM CHES, 10 mM Caps, 150 mM NaCl and 0.05% DDM, previously adjusted to the desired pH. For pKa determination the obtained spectra were normalized to the protein peak of 280 nm and the absorption maxima were plotted against pH.

For illumination experiments, solutions of *Ps*FAR-DM were prepared to have an optical density (OD) of ∼0.5 at an optical pathway of 10 mm. The dark-state spectrum was recorded then the main absorption band was illuminated using 490 nm LED (1W, Thorlabs).

### Ultrafast transient absorption spectroscopy

Femtosecond transient UV/vis absorption spectroscopy experiments were performed using a home-build setup described elsewhere^30^. In brief, a Ti:Sa chirped pulse regenerative amplifier (MXR-CPA-iSeries, Clark-MXR Inc.) was used as an fs-laser source, providing a pulse centered at 775 nm with a pulse width of 150 fs at a repetition rate of 1 kHz. Excitation pulses were generated using a two-stage NOPA (noncollinear optical parametric amplifier) setup with a prism compressor set between the NOPAs for pulse compression. The white light continuum probe pulses were generated by focusing the laser fundamental into a 5 mm thick CaF_2_-crystal which was continuously moved perpendicular to the incident beam. The generated white light pulses were split into a probe and a reference beam. The reference was guided directly into a spectrograph. The probe was focused with the excitation pulse, directly into the sample while the probe collected afterwards and guided into another spectrograph. The spectrographs (AMKO Multimode) contained gratings with 1200 grooves/mm blazed at 500 nm and a photodiode array with a detection range set to 350 – 700 nm. The measurements were performed under magic angle conditions (54.7° difference in pump-probe polarization) to exclude anisotropic signal contributions. The samples were held in a fused silica cuvette using an optical path length of 1 mm and sample concentration was adjusted to an OD of ∼0.4 at the excitation wavelength.

### Transient flash photolysis spectroscopy

The pump pulse was generated using a Nd:YAG (Surelite I-20, Continuum) operated at 20 Hz. The generated THG at 355 nm was used to pump an optical parametric oscillator (OPO) (preciScan, GWU Lasertechnik). The monochromatic probe light was generated by a xenon Lamp (LC-08, Hamamatsu) and wavelength selection was performed using two monochromators, one before and one after the probe light had passed the sample. The detection system consists of a photomultiplier tube and home-build amplification electronics, mounted directly after the second monochromator. The signal was digitized by two oscilloscopes (5244B and 5244D, Pico Technology, U.S.A.). Therefore, signals were detected every 56 ns up to 50 ms and every 10 µs for up to 10 s. After merging both datasets, the traces were averaged using a combined linear and logarithmic timescale to obtain feasible data sizes.

Sample measurements were performed using a 2 mm x 10 mm quartz glass cuvette with the probe light passing the sample with an optical path of 10 mm while the pump passed the sample perpendicular to the probe with an optical path length of 2 mm. The absorbance changes were monitored from 300 nm to 660 nm in 10 nm steps. Each wavelength was averaged using at least 30 scans to improve S/N. The excitation wavelength was set to 510 nm using an average pulse energy of ∼3.2 mJ/cm^2^.

### Analysis of time-resolved spectroscopic data

Data analysis was performed using OPTIMUS (http://www.optimusfit.org/)^31^. We applied a sequential model to perform global target analysis (GTA) on all time-resolved datasets. The analysis fits the model directly to the experimental data and results in the lifetimes, the decay-associated spectra (DAS) and the evolution-associated spectra (EADS) for each kinetic component. In contrast, the EADS contain pure spectral information while the DAS highlight the decay and rise of positive and negative bands associated with signal evolution.

### Cryo-EM grid preparation and data collection

All samples were concentrated to 60 mg/ml using 100,000 MWCO concentrators (Millipore) at pH 8.0 and later diluted with a buffer containing 20 mM Tris-HCl, pH 8.0, 100 mM NaCl and 0.04% DDM to the final concentration of 7 mg/ml. Then, 3 µl of the sample were applied onto freshly glow-discharged (30s at 5 mA) Quantifoil grids (Au R1.2/1.3, 300 mesh) at 20□°C and 100% humidity and plunged-frozen in liquid ethane. The cryo-EM data were collected using 300□keV Krios microscope (Thermo Fisher), equipped with Gatan K3 detector.

### Cryo-EM data processing

All steps of data processing were performed using cryoSPARC v.4.0.2^32^ (Fig. S2). Motion correction and contrast transfer function (CTF) estimation were performed with default settings for all datasets. Particle picking was performed using Topaz^33^ pre-trained model, followed by duplicate removal with 50 Å distance. Picked particles were extracted with 3-5x binning (up to 128 px). An initial set of particles was cleaned using two rounds of 2D classification (first round: 80 classes, 80 iterations, batch size 200-400, use clamp-solvent: true; second round: 20-40 classes, 40 iterations, batch size 50-200, use clamp-solvent: true). After that, particles were cleaned using a “3D classification” (*ab initio* model generation with 3 classes, followed by heterogeneous refinement). These particles were re-extracted without binning, followed by homogeneous refinement (C5 symmetry, with per-particle CTF and defocus refinement) and local refinement (C5 symmetry), yielding final maps.

### Model building and refinement

The pentameric models of *Ps*PR and *Ps*FAR were generated using Alphafold3^34^ and docked as a rigid body into cryo-EM maps manually in Chimera. Further refinement was performed using Refmac^35^, Phenix^36,37^ and Coot^38^, producing the final statistics described in Table S1. Visualization and structure interpretation were carried out in UCSF Chimera^39,40^ and PyMol (Schrödinger, LLC).

