## Supplementary Materials for "Structural basis for no retinal binding in flotillin-associated rhodopsins"

**Table S1. Cryo-EM data collection, refinement, and validation.**

| **Protein** | *Ps*FAR | *Ps*PR | *Ps*FAR-DM |
| --- | --- | --- | --- |
| **PDB ID** | 9R21 | 9R22 | 9R23 |
| **Data collection and processing** |  |  |  |
| Voltage (kV) | 300 | 300 | 300 |
| Electron exposure (e^−^/Å^2^) | 50 | 50 | 50 |
| Defocus range (μm) | −0.5 to −2.0 | −0.5 to −2.0 | −0.5 to −2.0 |
| Pixel size (Å) | 0.836 | 0.836 | 0.836 |
| Micrographs collected | 4478 | 6112 | 2015 |
| Micrographs processed | 3792 | 4589 | 1678 |
| Symmetry imposed | C5 | C5 | C5 |
| Initial dataset (# of particles) | 619,824 | 2,107,133 | 687,836 |
| Final dataset (# of particles) | 212,403 | 497,291 | 116,149 |
| Map resolution (Å) FSC_01.43_ | 2.56 | 2.48 | 2.81 |
| **Refinement** |  |  |  |
| Initial model used | Alphafold3 | Alphafold3 | *Ps*FAR (present work) |
| Map-sharpening *B* factor (Å^2^) | 123.7 | 121.8 | 131 |
| No. atoms |  |  |  |
| Protein | 9,230 | 8,895 | 9,194 |
| Retinal | - | 100 | 100 |
| Lipid | 904 | 543 | 700 |
| Water | 97 | 105 | 45 |
| *B*-factors (Å^2^) |  |  |  |
| Protein | 39.8 | 41.6 | 48.2 |
| Retinal | - | 31.8 | 43.0 |
| Lipid | 94.2 | 88.0 | 113.2 |
| Water | 42.3 | 39.4 | 38.7 |
| R.m.s. deviations |  |  |  |
| Bond lengths (Å) | 2.120 | 0.013 | 0.010 |
| Bond angles (°) | 0.011 | 1.941 | 1.788 |
| **Validation** |  |  |  |
| MolProbity score | 2.19 | 1.73 | 1.87 |
| Clash score | 11.88 | 9.09 | 9.33 |
| Poor rotamers (%) | 4.58 | 2.11 | 3.16 |
| Ramachandran plot (%) |  |  |  |
| Favored | 97.52 | 99.02 | 98.61 |
| Allowed | 2.48 | 0.98 | 1.39 |
| Outliers | 0 | 0 | 0 |
| Model to map fit CC | 0.90 | 0.88 | 0.86 |

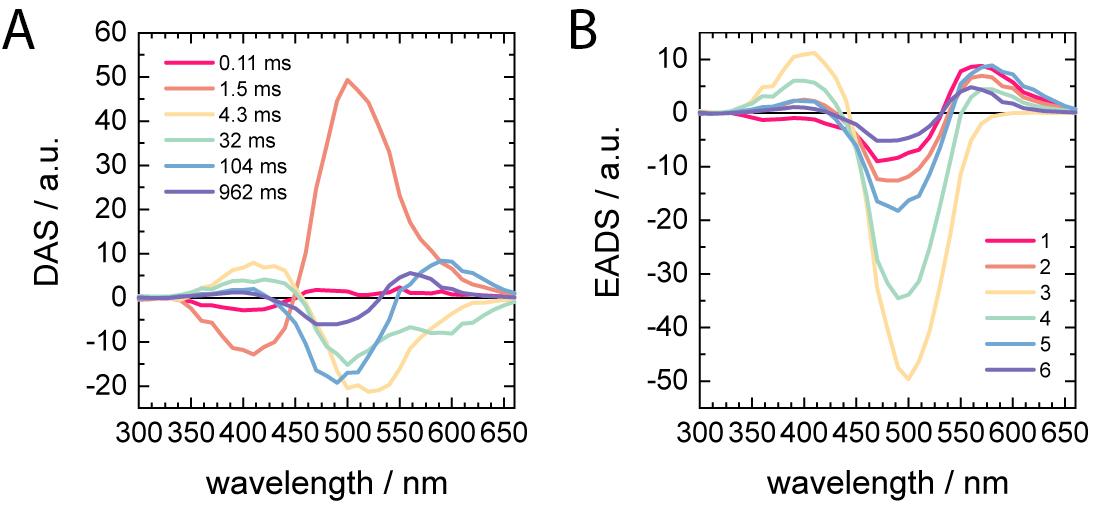

**Fig. S1. Kinetic analysis of ms-dynamics of *Ps*PR obtained by global target analysis using a sequential model. A.** Corresponding decay-associated spectra (DAS) highlighting the spectral changes associated with each of the six lifetimes. **B.** Evolution-associated difference spectra (EADS) displaying the spectral composition of each intermediate state. Spectral and kinetic decomposition allows the assignment of the components to construct the photocycle model following the spectral position and nomenclature originally derived for bacteriorhodopsin^13^ shown in Fig. 1E. The first two lifetimes of 0.11 ms and 1.5 ms describe the decay of the K intermediate towards the M state, spectrally characterized by a blue shift originating in the deprotonation of the RSB. M proceeds with a lifetime of 4.3 ms towards N, shifting the absorption towards the bleach. The N state decays with 32 ms and forms the red-shifted O_1_ intermediate. O_1_ proceeds to O_2_ with a lifetime of 104 ms before the decay of O_2_ recovers the ground state with a lifetime of 962 ms.

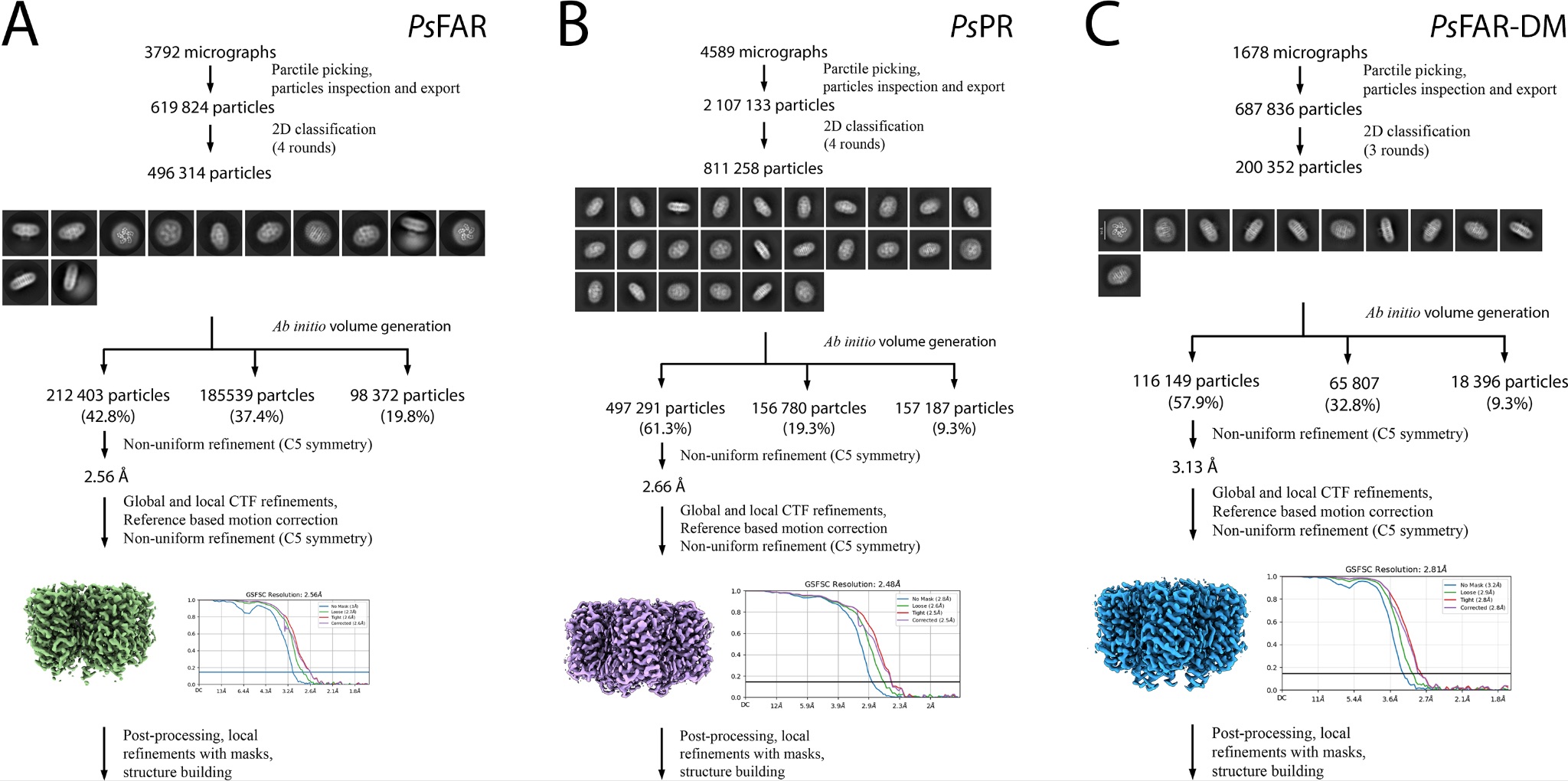

**Fig. S2. Workflow for solving the structures using cryoSPARC. A.** *Ps*FAR. **B.** *Ps*PR. **C.** *Ps*FAR-DM.

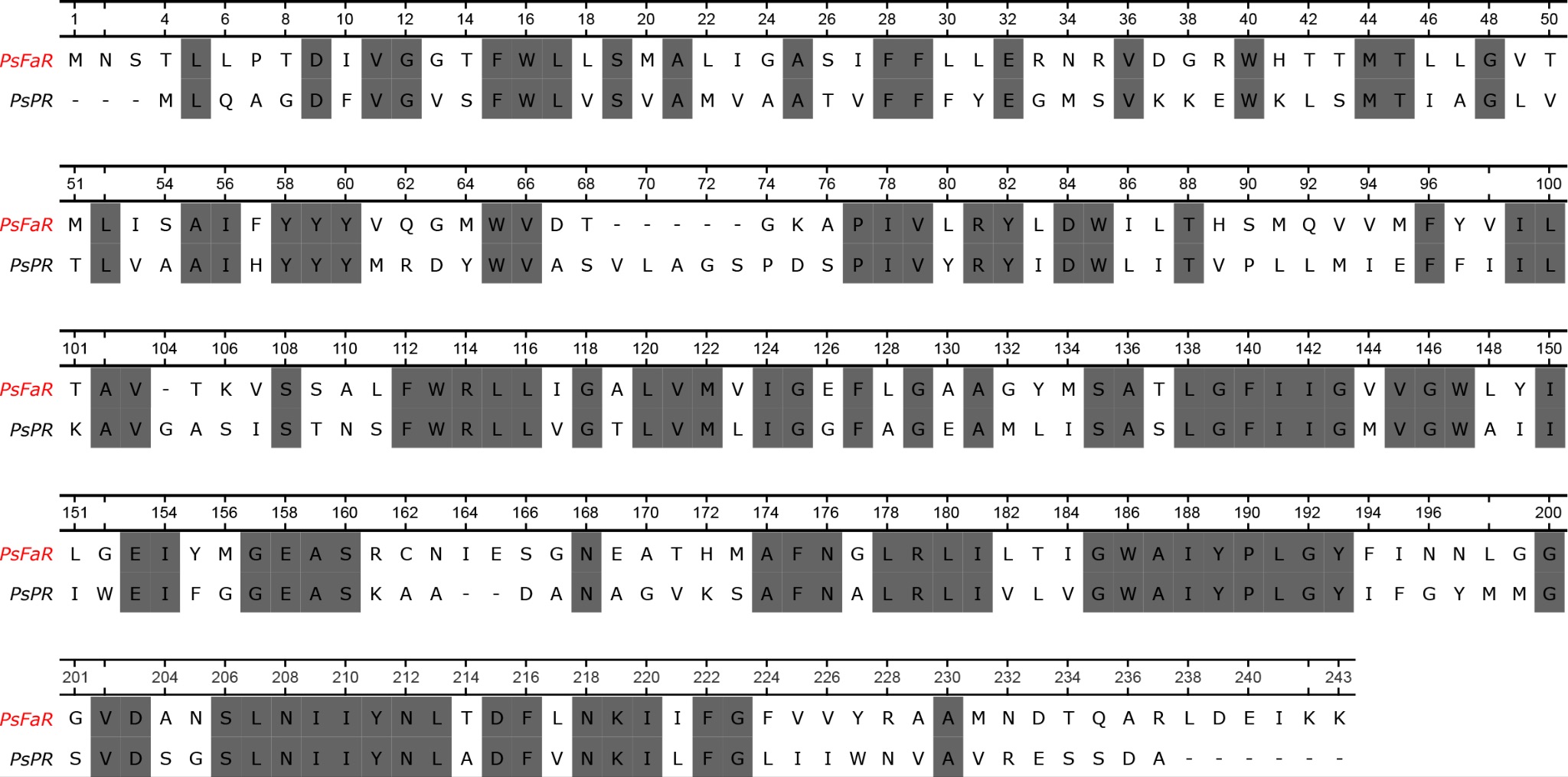

**Fig. S3. Sequence alignment of *Ps*FAR and *Ps*PR.**

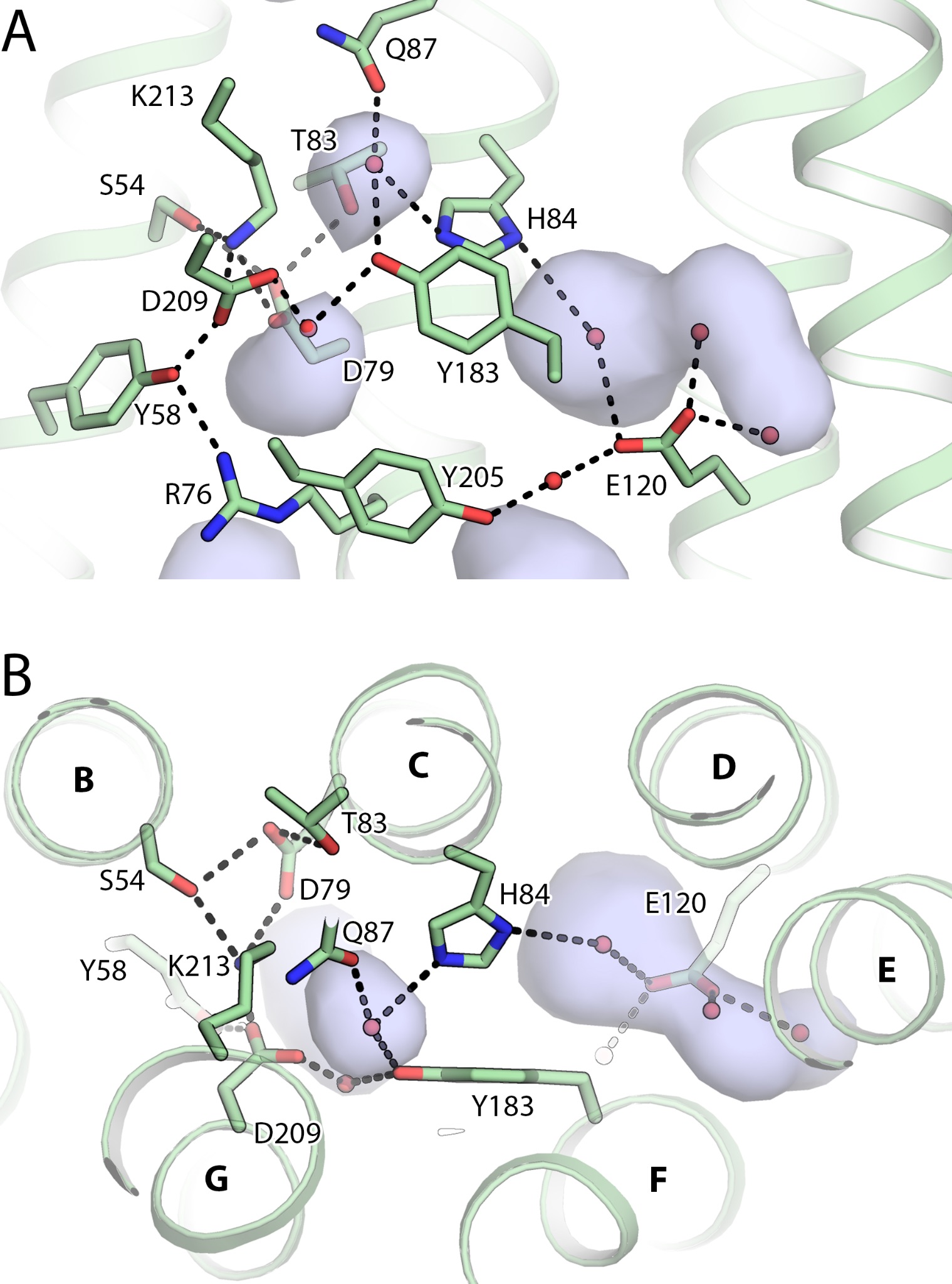

**Fig. S4. Hydrogen bond network in the central region of *Ps*FAR. A.** Side view. Helices F and G are not shown for clarity. **B.** Section view from the cytoplasmic side.

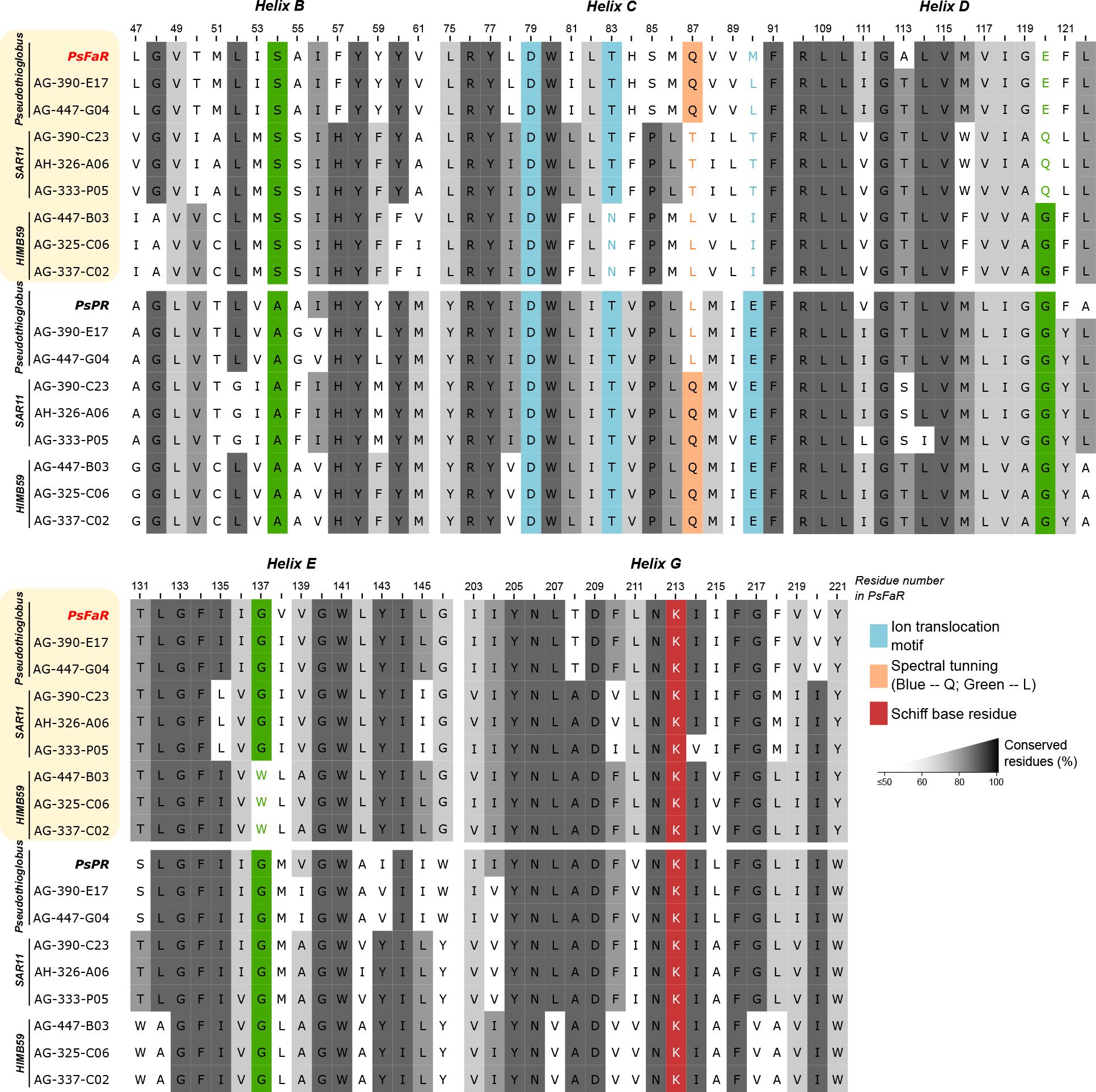

**Fig. S5. Sequence alignment of representative paralog FARs and PRs.** Position of S54 in *Ps*FAR is indicated in green to highlight conservativity of the residue within FARs but not PRs.

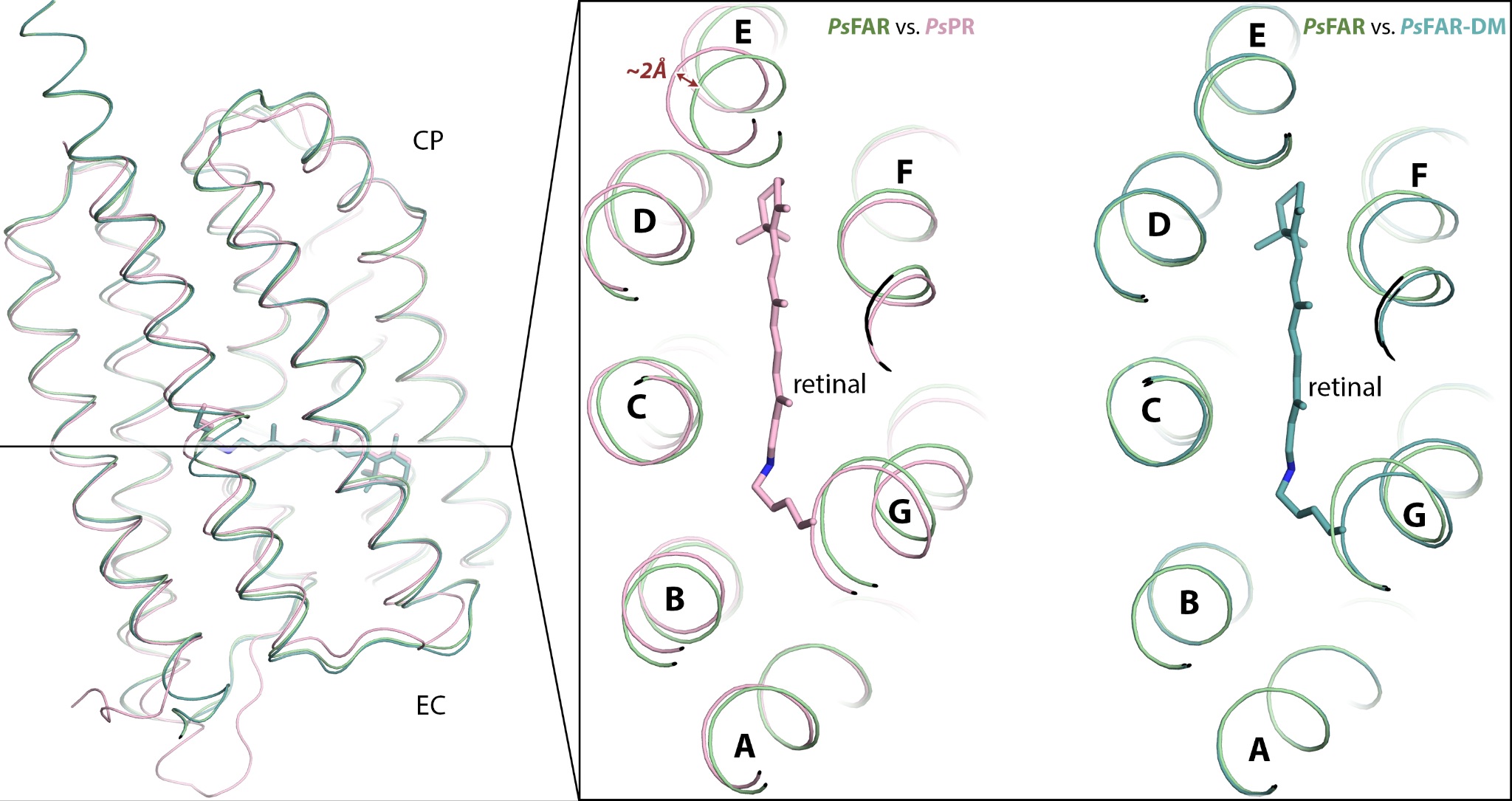

**Fig. S6. Comparison of helices positions in the central region of *Ps*FAR, *Ps*PR, and *Ps*FAR-DM.** Side view (left panel) and section view (right panel) of the structural alignment of the *Ps*FAR (green), *Ps*PR (pink), and *Ps*FAR-DM (deep cyan) protomers. The most pronounced shift of helix E in *Ps*PR and *Ps*FAR is indicated with a red arrow.

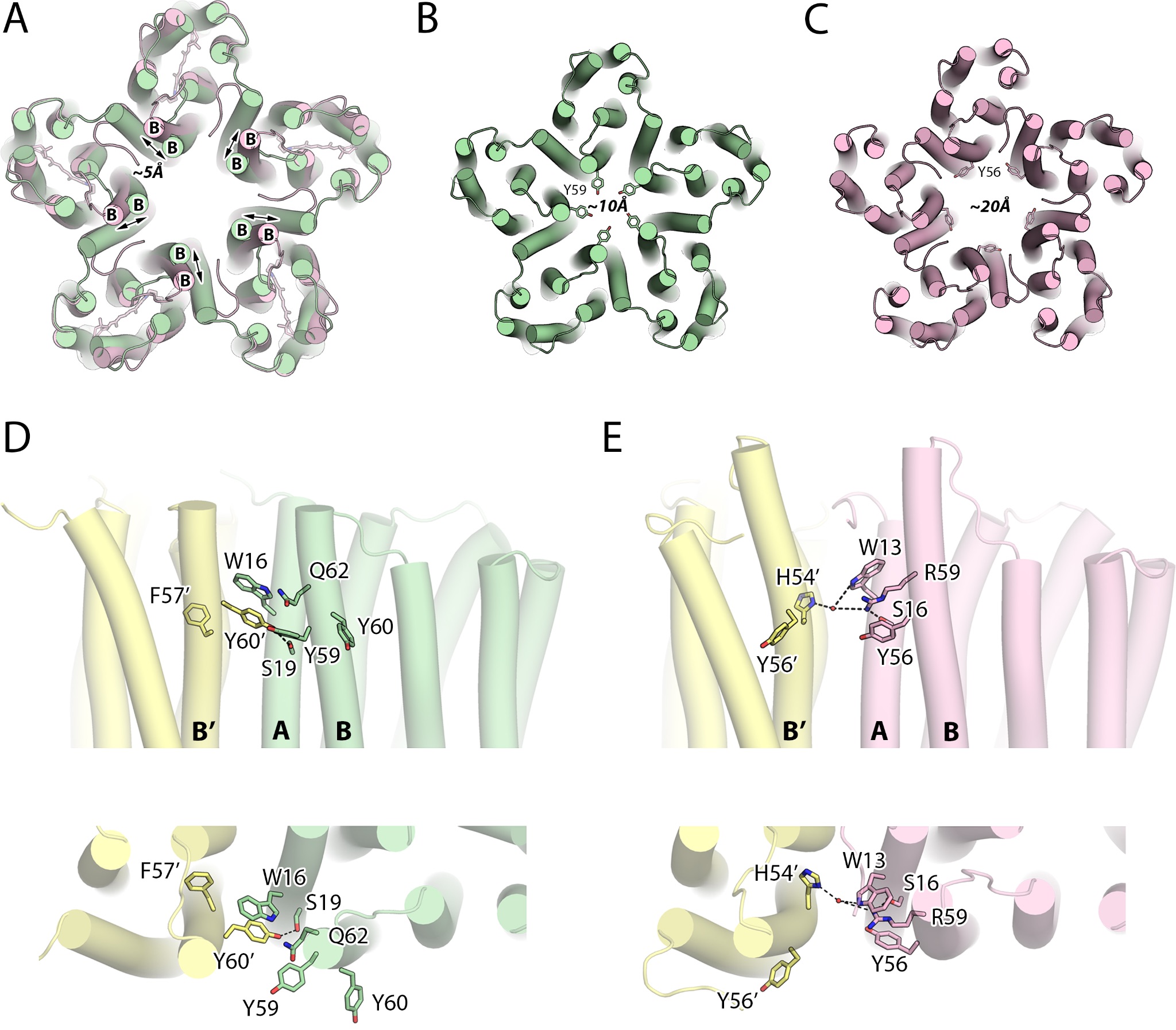

**Fig. S7. Comparison of interprotomeric contacts in *Ps*FAR and *Ps*PR. A.** View from the extracellular side on the structural alignment of *Ps*FAR and *Ps*PR. The difference in the tip of helix B position is indicated with black arrow. **B.** Y59 arrangement in *Ps*FAR. **C.** Y56 arrangement in *Ps*PR. The approximate diameters of the narrowest section of the central region are given in Å. **D.** Detailed view of the interprotomeric contacts at the extracellular side of the *Ps*FAR pentamer. Views from the central part of the pentamer (top panel) and from the extracellular side (bottom panel) are shown. **E.** Detailed view of the interprotomeric contacts at the extracellular side of the *Ps*PR pentamer. Views from the central part of the pentamer (top panel) and from the extracellular side (bottom panel) are shown. Hydrogen bonds mediating the contacts are shown with black dashed lines. Neighboring protomer is colored yellow.

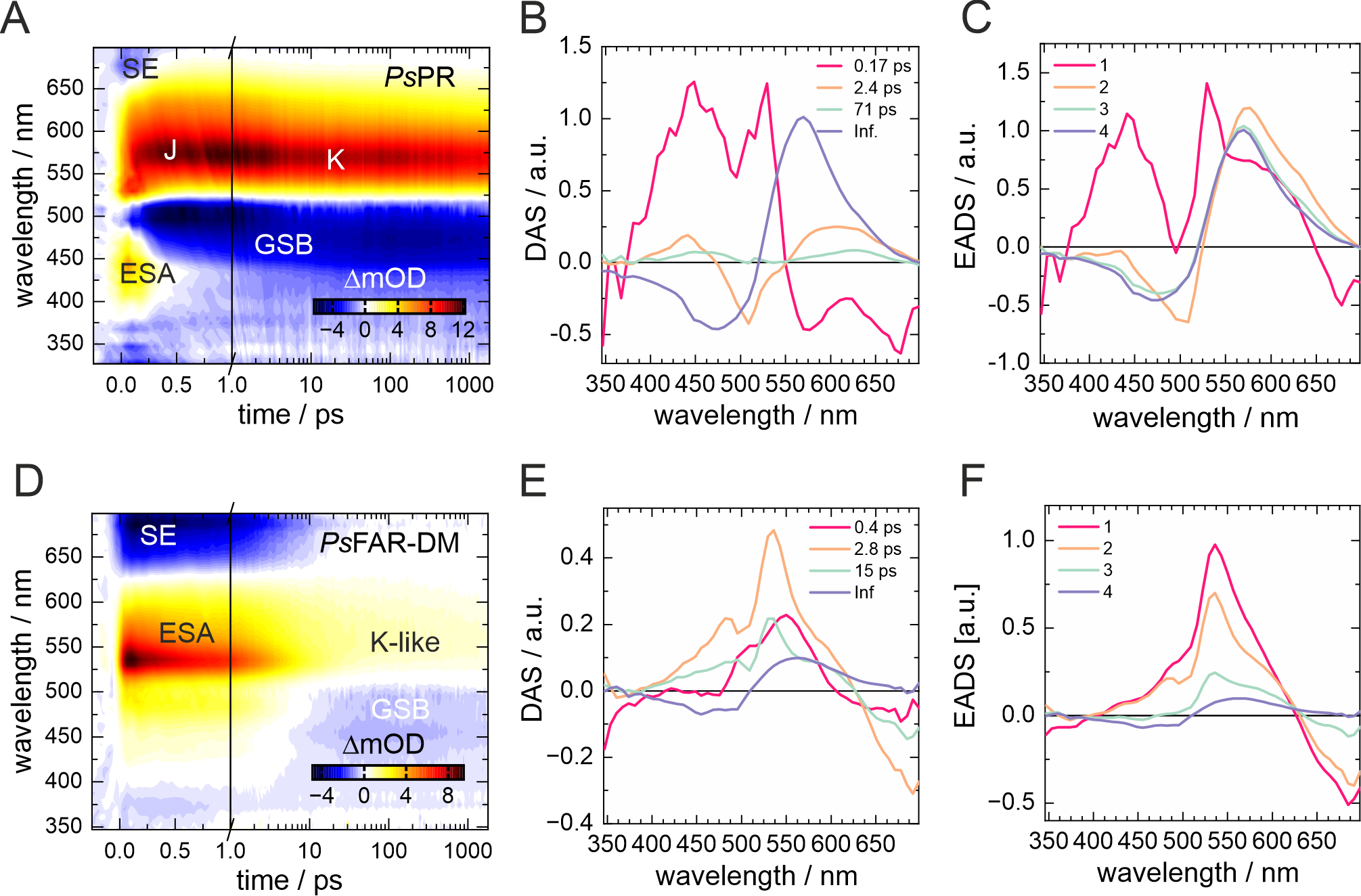

**Fig. S8. Ultrafast spectroscopy of *Ps*PR and *Ps*FAR-DM. A.** 2D-contour plot of a fs-TA measurement of *Ps*PR (pH 9.1) excited at 510 nm. **B.** Corresponding decay-associated spectra (DAS) and **C.** evolution-associated difference spectra (EADS) obtained by global target analysis of *Ps*PR **D.** 2D-contour plot of a fs-TA measurement of *Ps*FAR-DM (pH 8.0) excited at 490 nm. **E.** DAS and **F.** EADS obtained by global target analysis of *Ps*FAR-DM. The timescale of the transient maps is linear until 1 ps and logarithmic afterwards. In the dynamics of *Ps*PR the first lifetime of 170 fs is assigned to the excited state decay and the formation of the J intermediate. Within 2.4 ps J decays forming the K intermediate, which persists beyond the observation limit of 1.8 ns. In contrast, *Ps*FAR-DM required three time constants of 0.4 ps, 2.8 ps and 15 ps to describe the excited state decay adequately. The decay of the excited state results in the formation of a K-like intermediate, outlasting the observed timescale.
